# A human factor H-binding protein of *Bartonella bacilliformis* and potential role in serum resistance

**DOI:** 10.1101/2021.04.13.439661

**Authors:** Linda D. Hicks, Shaun Wachter, Benjamin J. Mason, Pablo Marin Garrido, Mason Derendinger, Kyle Shifflett, Michael F. Minnick

## Abstract

*Bartonella bacilliformis* is a Gram-negative bacterium and etiologic agent of Carrión’s disease; a potentially life-threatening illness endemic to South America. *B. bacilliformis* is a facultative parasite that infects human erythrocytes (hemotrophism) and the circulatory system, culminating in a variety of symptoms, including a precipitous drop in hematocrit, angiomatous lesions of the skin (verruga peruana) and persistent bacteremia. Because of its specialized niche, serum complement imposes a continual selective pressure on the pathogen. In this study, we demonstrated the marked serum-resistance phenotype of *B. bacilliformis*, the role of factor H in serum complement resistance, and binding of host factor H to four membrane-associated polypeptides of ∼131, 119, 60 and 43 kDa by far-western (FW) blots. The ∼119-kDa protein was identified as ABM44634.1 by mass spectrometry; a protein annotated as a 116.5-kDa outer membrane autotransporter (encoded by the BARBAKC583_1133 locus). We designated the protein as factor <u>H</u>-<u>b</u>inding <u>p</u>rotein <u>A</u> (FhbpA). FhbpA possesses three structural motifs common to all autotransporter proteins (i.e., a signal peptide, autotransporter β-barrel domain and passenger domain). Recombinant FhbpA passenger domain, but not the recombinant autotransporter domain, was able to bind human factor H when analyzed by FW blots. Phylogenetic analyses of the passenger domain suggest that it is well-conserved among *Bartonella* autotransporters, with closest matches from *Bartonella schoenbuchensis*. Transcriptomic analyses of *B. bacilliformis* subjected to conditions mimicking the sand fly vector or human host, and infection of human blood or vascular endothelial cells showed maximal expression of *fhbpA* under human-like conditions and during infection of blood and endothelial cells. Expression during HUVEC infection was significantly higher compared to all other conditions by DESeq2. Surface binding of serum factor H by FhbpA is hypothesized to play a protective role against the alternative pathway of complement fixation during *B. bacilliformis* infection of the human host.

**Author Summary:** *B. bacilliformis* is a bacterial pathogen that colonizes the circulatory system of humans, where it can cause a life-threatening illness unless treated. Serum complement is a major effector of innate humoral immunity and a significant obstacle that must be evaded for successful survival and colonization by pathogens, especially those residing in the vasculature. In this study, we examined the serum complement resistance phenotype of *B. bacilliformis* and identified four membrane-associated proteins that bind serum factor H; a protein used by the host to protect its own tissues from complement activation. One of the proteins was identified by mass spectrometry, characterized, and designated factor <u>H</u>-<u>b</u>inding <u>p</u>rotein <u>A</u> (FhbpA). FhbpA is a predicted autotransporter, and we determined that the translocated “ passenger” domain of the protein is responsible for binding factor H. We also determined that expression of the *fhbpA* gene was highest during infection of human blood and especially vascular endothelial cells or under conditions that simulate the human host. The results suggest that FhbpA binding of host serum factor H protects the bacterium against complement activation during infection.

## Introduction

*Bartonella* are arthropod-transmitted, Gram-negative bacteria that parasitize the circulatory system of mammals, spanning the gamut of rodents to cetaceans. Three pathogenic species cause most infections of humans, including *Bartonella bacilliformis, Bartonella quintana* and *Bartonella henselae*; the agents of Carrión’s disease, trench fever, and cat-scratch disease, respectively. Bartonelloses present with a wide range of symptoms and syndromes, such as chronic asymptomatic bacteremia, malaise, fever, myalgia, bacillary angiomatosis, bacillary peliosis, infectious endocarditis and hemolytic anemia. Nevertheless, conserved attributes of *Bartonella*’s pathogenesis form a foundation for these various manifestations. First, all bartonellae are hemotrophic; i.e., they infect erythrocytes, presumably to fulfill their extraordinary requirement for heme [1]. Hemotrophy is a highly unusual parasitic strategy for bacteria, and it contributes to the severe hemolytic anemia during the acute (hematic) phase of Carrión’s disease and the persistent bacteremia common to all types of bartonelloses [1]. Second, *Bartonella*’s infection of vascular endothelial cells can provoke pathological angiogenesis in humans, culminating in bacillary angiomatosis (*B. quintana* or *B. henselae*), verruga peruana (*B. bacilliformis*), or bacillary peliosis of the liver or spleen (*B. henselae*) [1].

The complement system of vertebrates consists of over thirty proteins synthesized by the liver and released into serum, where they influence both cellular and inflammatory processes in the humoral immune compartment. Complement activities include opsonization (by C3b), activating the discharge of pre-formed inflammatory mediators of granulocytes such as histamine release during mast cell degranulation (by the anaphylatoxins C3a, C4a, C5a), leukocyte chemotaxis and recruitment to an area of microbial challenge (by C5a), and lysis of invading microbes (by the membrane attack complex, C5bC6C7C8C9). Complement activation proceeds by cascade-mediated processes initiated by: 1) IgG or IgM binding to an antigen (classical pathway), 2) C3b factor binding to an activator surface such as peptidoglycan or LPS (alternative pathway), and 3) binding of mannose-binding lectin (MBL) to a surface containing mannose (lectin pathway). While the classical pathway is initiated by IgG or IgM immunoglobulin binding to an antigen and therefore dovetails with the adaptive immune response, the alternative and lectin pathways are entirely innate forms of immunity and provide a first line of defense against microbial challenge.

Many bacterial pathogens of vertebrates have evolved mechanisms that confer resistance to serum complement in order to colonize the host. Serum resistance in bacteria is conferred by various molecular means, including capsular polysaccharides and lipopolysaccharide [2, 3], utilization of surface receptors that bind host serum factor H to accelerate the decay of C3 convertase (C3bBb), inactivation of C3b in collaboration with factor I of the alternative pathway [4, 5], prevention of IgM binding to inhibit activation of the classical pathway [6], binding of host C1 esterase inhibitor [7], and binding of the C4b-binding protein inhibitor of classical and lectin pathways [8]. In the present study, we describe a human serum factor H-binding protein of *B. bacilliformis* that is annotated as an autotransporter (ABM44634.1). To our knowledge, this is the first report describing a potential complement-resistance factor of *B. bacilliformis*.

## Materials and methods

### Ethics statement

The Institutional Biosafety Committee and Institutional Review Board at the University of Montana granted approval for experimental use of human blood (IBC 2019-05; IRB 120-20). Formal consent was obtained in verbal form from the blood donor (co-author MM).

### Bacterial strains and cultures

*B. bacilliformis* type strain KC583 (ATCC 35685; American Type Culture Collection; Manassas, VA) was used in all experiments. Cultivation of the bacterium was limited to six passages beyond the ATCC stock. Cultures were grown 4 d (approx. mid-log phase) at 30°C and 100% relative humidity on agar plates consisting of a heart infusion broth base (HIB; Becton Dickinson; Franklin Lakes, NJ) supplemented with 4% sheep erythrocytes and 2% filter-sterile sheep serum by volume (HIBB medium). Sheep erythrocytes and sera were purchased from Quad Five; Ryegate, MT. *Escherichia coli* strains (**Table S1**) were grown in lysogeny broth (LB) or LB agar plates for 16 h at 37°C. Antibiotic supplements were added to media as needed for selection (e.g., kanamycin 50 μg/ml, ampicillin 100 μg/ml).

### Serum resistance assays

For serum resistance assays, four 4-d-old *B. bacilliformis* cultures were harvested from HIBB plates into HIB at 25°C. The cell suspension was gently vortexed, centrifuged for 1 min at 16,100 x g and the supernatant discarded. The remaining pellet was gently and thoroughly resuspended into 1 ml HIB. From this suspension, 100 μl aliquots (in duplicate) were removed and added to: 1) pooled human serum complement (HSC; Innovative Research; Novi, MI) in HIB to give a 50% serum concentration, 2) pooled human serum complement inactivated by treating for 30 min at 56°C then diluted in HIB to give a 50% concentration (iHSC), or 3) human factor H-depleted serum (FHDS; Complement Technology; Tyler, TX) diluted in HIB to give a 50% concentration. The resulting cell mixtures were incubated for 1h at 30°C with gentle vortexing by hand at 5-min intervals. Following incubation, the mixtures were placed on ice and immediately ten-fold serially diluted to 10^−6^ in ice-cold HIB. Aliquots of the dilutions (100 μl) were plated onto HIBB in duplicate and incubated for 7d at 30°C. Average CFU’s were determined by manual colony counting.

### Sodium dodecyl sulfate-polyacrylamide gel electrophoresis (SDS-PAGE) and far-western (FW) blots

Whole-cell lysates of *B. bacilliformis* were prepared by harvesting 4-d-old cultures on HIBB plates into ice-cold HIB. The cell suspension was centrifuged for 5 min (6000 x g, 4°C) and the pellet resuspended in 1 ml cold PBS (pH 7.4). After re-centrifuging, the final pellet was suspended in 1 ml PBS and frozen (−80°C) until used. Total membranes of *B. bacilliformis* were prepared as we previously described [9]. Protein concentrations were determined using a Pierce BCA Protein Assay kit as instructed by the manufacturer (Thermo Fisher; Waltham, MA).

Protein profiles were analyzed by SDS-PAGE using pre-cast 4-20% acrylamide gradient gels (Novex WedgeWell, Tris-glycine gel; Thermo Fisher) and 20 μg protein per well. Samples were solubilized in Laemmli 6X sample buffer, boiled 10 min and centrifuged 1 min (16,100 x g, 25°C) prior to loading the resulting supernatants onto gels. Protein banding patterns were visualized by staining gels with Coomassie brilliant blue R. Far-western (FW) blots were prepared by transferring proteins from un-fixed / unstained SDS-PAGE gels to supported nitrocellulose [0.45 μm pore; Cytiva, Marlborough, MA] [10], immediately following electrophoresis. FW blots were blocked overnight in PBS-T.3 [PBS (pH 7.4) and 0.3% Tween-20] containing 5% (w/v) non-fat dry milk. Blots were then probed for 60 min with human complement factor H (Complement Technology) at 5 ng/μl in PBS-T.3, followed by a 60-min incubation with mouse anti-human factor H antibodies (MilliporeSigma, St. Louis, MO) diluted 1:1000. After three 5-min washes in PBS-T.3, blots were re-probed with rabbit anti-mouse IgG peroxidase-conjugated antibodies (Bio-Rad / AbD Serotec; Hercules, CA) diluted 1:40,000 in PBS-T.3. Blots were then washed three times for 5 min in PBS-T.3 and developed with ECL reagents per manufacturer instructions (SuperSignal West Pico Chemiluminescent Substrate; Thermo Fisher). FW blots were visualized with a LAS-3000 digital imaging system (Fujifilm; Valhalla, NY).

### Mass spectrometry

Membrane-associated proteins of *B. bacilliformis* were prepared, separated by SDS-PAGE (20 μg protein per lane) and the resulting gels stained with Coomassie brilliant blue R, as above. Protein bands of interest were excised from gels and submitted to Alphalyse Laboratories (Palo Alto, CA), for mass spectrometry (MS). Briefly, samples were reduced and alkylated, then digested with trypsin. The resulting peptides were evaluated by matrix-assisted laser desorption / ionization tandem time-of-flight (MALDI-TOF/TOF) MS. For peptide fragmentation analysis (partial sequencing), MALDI MS/MS was employed. Database searches were done using the MS and MS/MS data and Mascot 2.4 software (Matrix Science; Boston, MA).

### Immunofluorescence analysis and UV microscopy

*B. bacilliformis* was grown 4 d on four HIBB plates and harvested into 2 ml HIB at 25°C. Cells were centrifuged for 5 min (6,000 x g, 25°C). The resulting pellet was washed 3 times in 2 ml PBS (pH 7.4, 25°C), with gentle vortexing and centrifugations, as above, after each wash. The final pellet was resuspended in 2 ml of 2% paraformaldehyde (in PBS) and incubated 45 min at 25°C to fix the cells. Fixing was quenched with 2 ml 0.1 M glycine (in PBS). Fixed bacteria were pelleted by centrifugation and washed 3 times with PBS containing 0.05% Tween-20 (PBS-T.05), as above. To block non-specific antibody binding, cells were resuspended in PBS containing 0.05% Tween 20 and 5% (v/v) donkey serum (PBS-TDS) and incubated 60 min at 25°C with rocking. This mixture was re-centrifuged, the pellet resuspended in 250 μl pooled HSC (Innovative Research) and then incubated for 30 min. Cells were then washed three times in PBS-TDS with centrifugations, as above, after each wash. The final pellet was resuspended in PBS-TDS and divided into 3 aliquots of equal volume in microcentrifuge tubes. Tubes were centrifuged and the pellets resuspended in: a) goat anti-human factor H antiserum (Complement Technology), b) PBS or c) pooled goat normal serum (Quad Five) diluted in PBS-TDS (1:100). Mixtures were incubated for 60 min at 25°C with gentle rocking, then re-centrifuged. Resulting pellets were washed 3 times with PBS-TDS with centrifugations between washes. The final pellets were resuspended in AlexaFluor 488 donkey anti-goat IgG (Thermo Fisher) diluted 1:100 in PBS-TDS, and incubated for 60 min at 25°C with gentle rocking. The cells were then washed 3 times in PBS-TDS with centrifugations after each wash, as above. The final pellet was resuspended in PBS-T.05, and wet mounts were prepared and observed with an Olympus BX51 phase contrast microscope equipped with a fluorescence illuminator (X-Cite 120Q; Excelitas Technologies; Waltham, MA), DP72 camera (Olympus; Center Valley, PA) and DP2-BSW acquisition software (Olympus).

### Cloning and expression of *fhbpA*

The Gateway system (Thermo Fisher) was used to clone predicted autotransporter and passenger domains of FhbpA (ABM44634.1). Briefly, synthetic oligonucleotides corresponding to 5’ and 3’ ends of the coding sequences for the domains (see **S1 Table** for details) were used to amplify respective targets by standard PCR. The PCR products were cloned into an entry vector using a pENTR/D-TOPO Cloning Kit, then transformed into *E. coli* (Top10) as instructed by the manufacturer (Thermo Fisher). Positive colonies were identified by PCR screening with the same primers used in cloning. Plasmids were purified from positive clones with a QIAprep Spin Miniprep plasmid kit as instructed (Qiagen; Germantown, MD), and verified by Sanger automated sequencing (ACGT; Germantown, MD). pENTR/D-TOPO insert DNA was transferred to a pET-DEST42 destination plasmid and used to transform *E. coli* BL21 Star (DE3) for expression using a Gateway LR Clonase II enzyme mix, as instructed by the manufacturer (Thermo Fisher). Positive clones were identified by PCR screening, and plasmid content was verified by Sanger automated sequencing, as above.

### Bioinformatic analysis

In order to determine those conditions that may regulate expression of BARBAKC583_1133, we analyzed *B. bacilliformis* whole transcriptome data [11] obtained from the Sequencing Read Archive database (accession number PRJNA647605) using DESeq2 software [12]. For the DESeq2 analysis, the p-value distribution of differentially expressed genes was re-calculated using the fdrtool package [13]. The resulting data more accurately reflected the desired null distribution of p-values and effectively made the analysis more stringent. TPM calculations were made using a python script located in a github repository (https://github.com/shawachter/TPM_Scripts).

### Software, graphics and statistics

The domains of FhbpA (ABM44634.1; BARBAKC583_1133 locus) were predicted using SignalP 4.1 [14] and SMART [15] for the secretory signal sequence, JPred4 [16] for a potential helical linker, and PROSITE-Expasy [17], BLAST [18] and SMART [15] for autotransporter (beta) and passenger domains. Structure predictions for FhbpA were modeled using Phyre2 [19]. Statistical analyses were done using Prism 9.0 software (GraphPad, La Jolla, CA) and student’s t-tests, where p values < 0.05 were considered significant. Phylogenetic analyses and trees were prepared using Mega 7.0 software [20]. Other figures and graphs were generated using PowerPoint and Excel software (Microsoft, Redmond, WA), respectively.

## Results

### Serum complement resistance of *B. bacilliformis* and involvement of factor H

The *B. bacilliformis* life cycle revolves around intracellular infection of human erythrocytes and vascular endothelial cells [21-23]. As such, complement imposes a persistent selective pressure, especially when the pathogen is extracellular. We were therefore curious about the serum-resistance of *B. bacilliformis*. To examine this phenotype, standard serum assays were done using commercially-available human serum components, while tailoring the assay for *Bartonella* (e.g., using HIB as a diluent). Results of the serum resistance assays showed that *B. bacilliformis* had a 63.3 ± 4.8% survival rate in human serum complement (HSC) at a 50% concentration, relative to untreated controls (**Fig. 1**). While this is a significant reduction compared to untreated controls (p<0.05), the results suggest that *B. bacilliformis* is decidedly resistant to complement. If human serum complement was inactivated at 56°C for 30 min (iHSC) and then used in assays at a 50% concentration, the percent survival was decreased to 80 ± 6.5%; a value that was not significantly different from the untreated controls, and implicated complement as the major bactericidal factor present in human serum (**Fig. 1**).

**Fig. 1.**
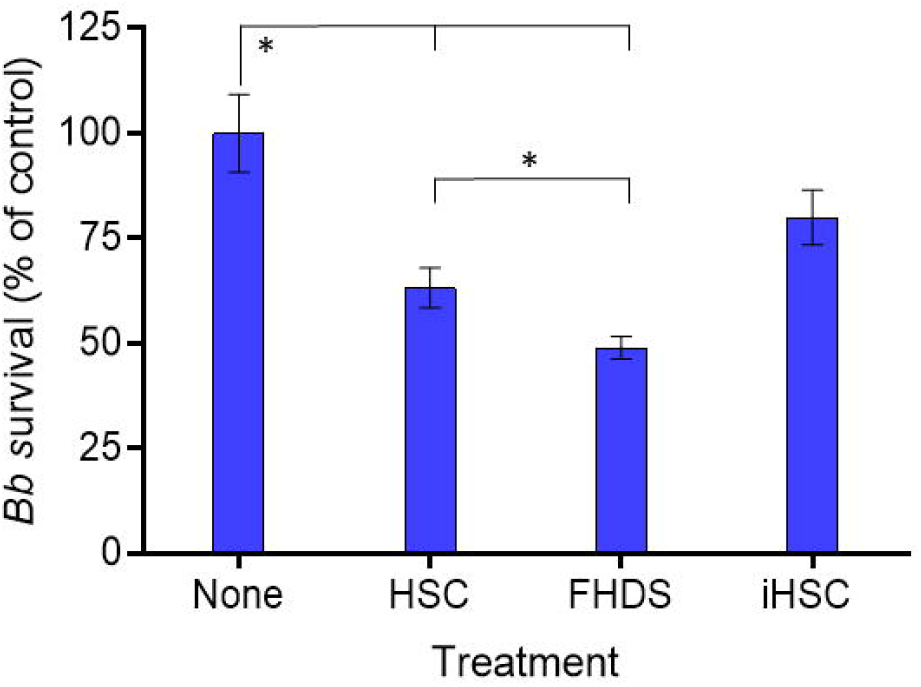
*B. bacilliformis* complement resistance and involvement of factor H. *B. bacilliformis* (strain KC583) had a mean survival rate of 63.3 ± 4.8% after a 1h incubation in pooled human serum complement (HSC, 50% concentration) vs. untreated controls. *B. bacilliformis* displayed a significantly lower survival rate (49 ± 2.7%) after a 1h incubation in factor H-depleted human serum (FHDS, 50% concentration) vs. those treated with HSC. The percent survival of untreated bacteria (None) and those treated with heat-inactivated human serum complement (iHSC) at a 50% concentration was not significantly different. Values represent the means of 4 independent serum assays ± SEM. Asterisks denote significant differences (p < 0.05) by unpaired student’s t tests.

To examine the role of factor H in conferring serum resistance to *B. bacilliformis*, assays were also done using human factor H-depleted serum (FHDS) at a 50% final concentration. Results of the FHDS assays showed a significant decrease in percent survival (−14.3%; p<0.05) compared to the HSC-treated bacteria, suggesting that serum factor H is used by *B. bacilliformis* to protect against complement activation via the alternative pathway (**Fig. 1**). Incomplete abrogation of resistance in the absence of factor H suggests that other types of complement resistance factors are employed by *B. bacilliformis*.

### Factor H binding to *B. bacilliformis* cells and proteins

Since factor H can bind to the surface of certain pathogenic bacteria and provides protection from complement, we tested for factor H binding to fixed, intact *B. bacilliformis* cells by imaging with immunofluorescence microscopy. Results of these experiments clearly showed a consistent and markedly greater intensity of fluorescence in bacteria probed with pooled HSC followed by goat anti-human factor H antiserum (**Fig. 2A**) relative to bacteria treated with pooled, naive goat serum or PBS in place of the factor H antiserum (**Figs. 2B, 2C**). These data suggest that factor H binds to the surface of intact *B. bacilliformis* cells.

**Fig. 2.**
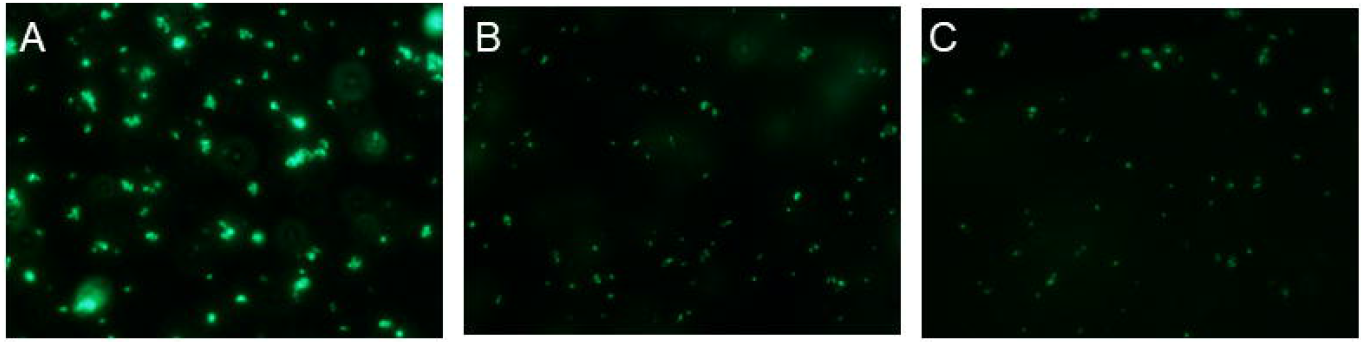
Human factor H binding to intact *B. bacilliformis* cells as demonstrated by immunofluorescence microscopy. Bacterial cultures were harvested at mid-log phase, fixed in paraformaldehyde, and quenched with glycine. Following three washes in PBS-T (PBS + 0.05% Tween 20), cells were incubated with pooled HSC for 30 min and washed 3 times in PBS-TDS (PBS-T + 5% pooled donkey serum). Mixtures were then incubated for 1 h with either: **A)** goat anti-human factor H antiserum, **B)** pooled goat serum, or **C)** PBS (pH 7.4). After 3 washes in PBS-TDS, factor H binding was visualized with AlexaFluor 488-conjugated donkey anti-goat IgG and fluorescence microscopy.

### Identity, phylogeny, and predicted structure of a factor H-binding protein of *B. bacilliformis*

To discover the surface-exposed, factor H-binding proteins (Fhbp’s) of *B. bacilliformis*, total membranes were purified from the bacterium, as we previously described [9]. The membrane-associated proteins were subsequently resolved by SDS-PAGE and blotted to nitrocellulose. The resulting blots were probed with human factor H to identify the membrane-associated Fhbp’s. FW blots consistently identified four prominent Fhbp’s of approximately 131, 119, 60 and 43 kDa that were enriched in the membrane fraction of the bacterium (**Fig. 3B, lane 2**). Of these, only the ∼60-kDa protein was detected in both the total cell lysate and membrane fractions of *B. bacilliformis*. Protein bands corresponding to the ∼131-kDa and ∼119-kDa polypeptides were submitted to Alphalyse, Inc., for analysis by MALDI-TOF/TOF mass spectrometry (MS). MS identified the ∼119-kDa polypeptide band as ABM44634 (GenBank); a protein annotated as an outer membrane autotransporter. This protein was subsequently designated factor <u>H</u>-<u>b</u>inding <u>p</u>rotein <u>A</u> (FhbpA).

**Fig. 3.**
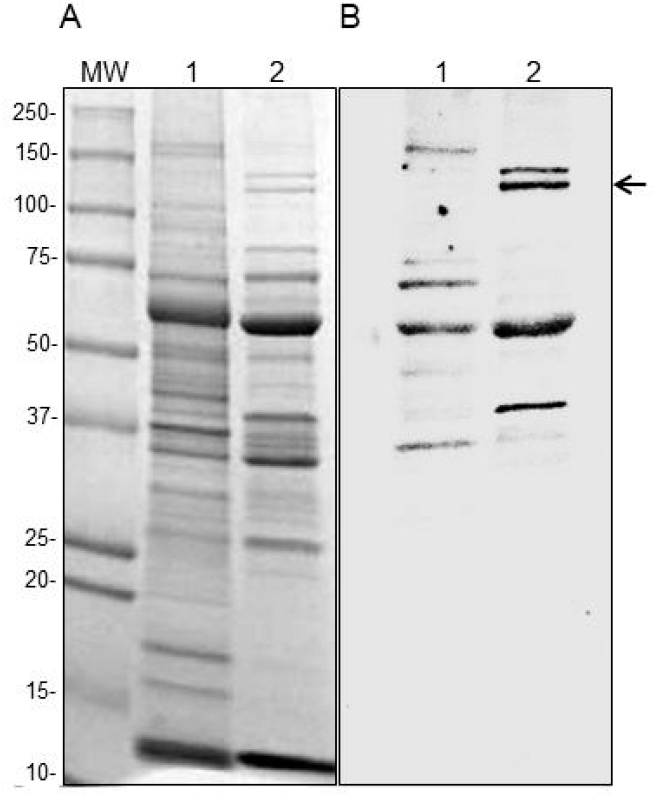
Four membrane-associated *B. bacilliformis* proteins bind human serum factor H. **A)** Coomassie blue-stained SDS-PAGE gel (4-20% w/v acrylamide gradient), containing: MW, protein molecular weight standards; lane 1, *B. bacilliformis* whole-cell lysate, and lane 2, a *B. bacilliformis* membrane preparation. **B)** Corresponding FW blot probed successively with: human factor H, mouse anti-human factor H antibody, and rabbit anti-mouse IgG::HRP. Four prominent *B. bacilliformis* factor-H binding proteins (Fhbp’s) of ∼131, 119, 60 and 43 kDa were enriched and identified in the membrane preparation (lane 2). The ∼119-kDa FhbpA protein band is arrowed. Molecular weights from protein standards in MW are shown to the left in kDa.

The predicted 1,058 amino acid sequence of FhbpA was used in BLAST searches to help elucidate the protein’s function and phylogeny. While the results of the searches strongly suggested that FhbpA was an autotransporter protein, use of the entire sequence as a search query resulted in homology mapping primarily to the protein’s predicted secretory signal sequence and/or autotransporter beta barrel domains (data not shown). We therefore confined the BLASTP search to the 720-residue, predicted passenger domain of FhbpA (amino acids 26-745) as the search query, since this is typically the “functional”, transported portion of autotransporter proteins. Results of this analysis showed that the twelve homologs with the highest total scores were all predicted autotransporter proteins of *Bartonella*, with passenger domains of *Bartonella schoenbuchensis* autotransporters most closely-related to that of FhbpA (**Fig. 4**).

**Fig. 4.**
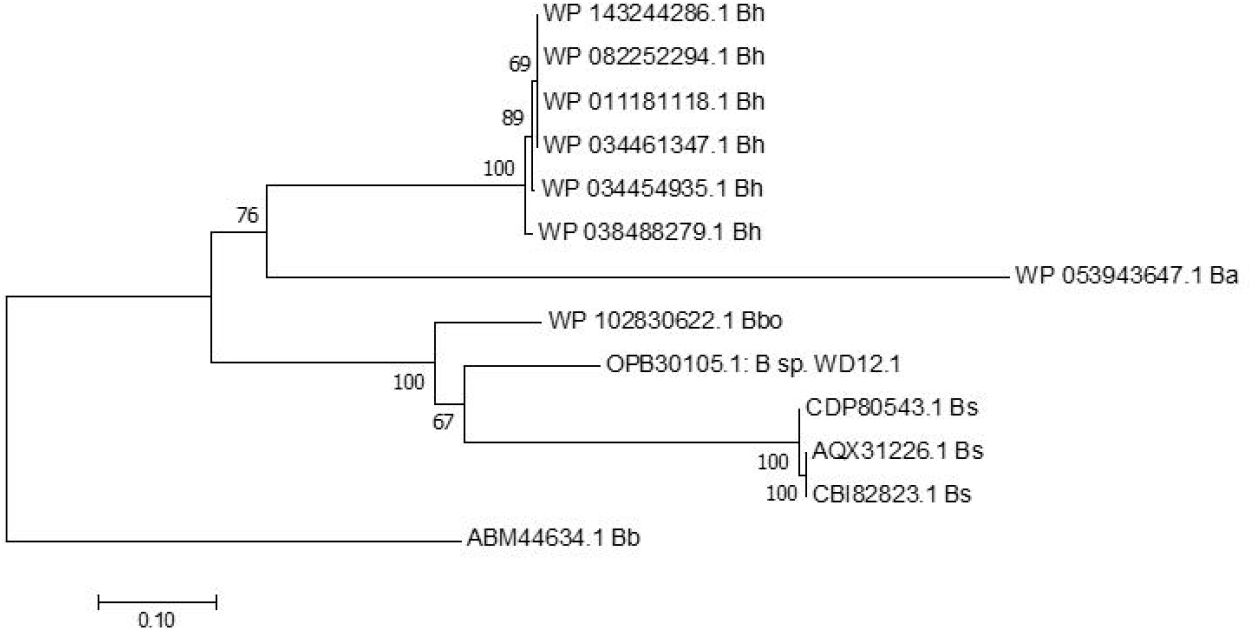
Phylogenetic analysis. Neighbor-joining tree drawn to scale for the twelve BLASTP hits with the highest total scores from other bacteria, using the predicted passenger domain of *B. bacilliformis* FhbpA (ABM44634.1; residues 26-745) as a search query. All homologs identified were predicted autotransporter proteins. Phylogenies were computed using the Poisson correction method. Bootstrap values (1,000 replicates) are shown at the nodes. Abbreviations: Ba, *B. ancashensis*; Bb, *B. bacilliformis*; Bbo, *B. bovis*; Bh, *B. henselae*; Bs, *B*. schoenbuchensis; and B sp. WD12.1, an undescribed *Bartonella* species.

Structure predictions of FhbpA were also done *in silico* using several programs available online (see Materials and methods). From these results, we determined the overall arrangement of the FhbpA protein precursor (**Fig. 5A**). The immature protein includes a 25-amino acid signal sequence, a 720-residue passenger domain, a 23-residue helical linker, a 9-residue spacer, the 268-residue autotransporter domain, and a 13-amino acid tail with a carboxy-terminal phenylalanine residue. FhbpA’s predicted passenger and autotransporter domains were also analyzed and modeled using Phyre2 [19]. These results showed that 430 residues (∼60% of the predicted passenger domain) could be modeled with 99.7% confidence by the *Bordetella pertussis* virulence factor p.69 pertactin (**Fig. 5B**). In addition, 239 residues of FhbpA (89% of the predicted autotransporter domain) could be modeled with 100.0% confidence by the pre-cleavage structure of the EspP autotransporter serine protease of *E. coli* O157:H7 (**Fig. 5C**).

**Fig. 5.**
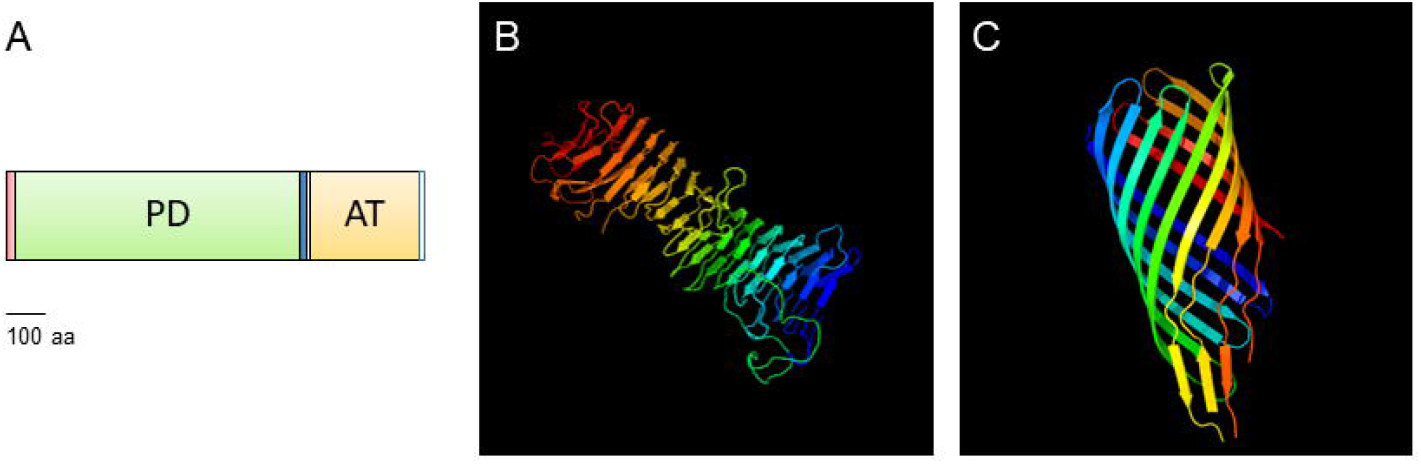
Structure predictions for FhbpA. **A)** Linear arrangement of the FhbpA precursor, including a 25-amino acid (aa) signal peptide (red), 720-aa passenger domain (PD, green), 23-aa linker (blue), 9-aa spacer, 268-aa autotransporter domain (AT, yellow) and a 13-aa tail. **B)** FhbpA passenger domain structure prediction by Phyre2 [19]. 430 residues (∼60% of the predicted domain) were modeled with 99.7% confidence by the single highest-scoring template; the *Bordetella pertussis* virulence factor p.69 pertactin. **C)** FhbpA autotransporter domain structure prediction by Phyre2 [19]. 239 residues (∼89% of the predicted domain) were modeled with 100.0% confidence by the single highest-scoring template; the pre-cleavage structure of the *E. coli* O157:H7 autotransporter serine protease, EspP.

### Identification of the factor H-binding domain of FhbpA

Passenger domains of several autotransporter proteins are involved in binding various host proteins, including serum factor H and extracellular matrix proteins (e.g., fibronectin and laminin). We therefore hypothesized that FhbpA’s factor H-binding activity likely involved its passenger domain. To address the hypothesis, we prepared FW blots with soluble and insoluble fractions of *E. coli* strains (see **S1 Table** for details) ectopically expressing either FhbpA’s autotransporter domain (strain LDH444) or passenger domain (strain LDH555). An *E. coli* strain expressing recombinant *B. bacilliformis* GroES (strain LDH333) was used as a negative control. FW blots probed with a Nickel-HRP probe for the His_6_ tag of each fusion protein detected three recombinant proteins (**Fig. 6B**), including the ∼15.8-kDa GroES band and autotransporter bands of ∼37-kDa (major) and ∼35.2-kDa (minor) (predicted molecular mass of ∼31.1 kDa) in the insoluble fractions of LDH333 and LDH444, respectively (**Fig. 6B**; **lanes 2 and 4**). Surprisingly, passenger domain protein bands of approximately 82.2, 70, 63.5, 45.7, and 31.9 kDa were all detected by the nickel-HRP probe in the insoluble fraction of strain LDH555 (**Fig. 6B**; **lane 6**). Protein bands of the same molecular weight could also be seen in the Coomassie blue-stained gel of strain LDH555 (**Fig. 6A; lane 6**). The 82.2-kDa protein band is presumably the full-length FhbpA passenger domain, with a predicted molecular mass of ∼82.8 kDa, whereas the proteins of lower molecular weight are possibly partial-proteolysis fragments with intact His_6_ tags on their C termini that are recognized and bound by the nickel-HRP probe.

**Fig. 6.**
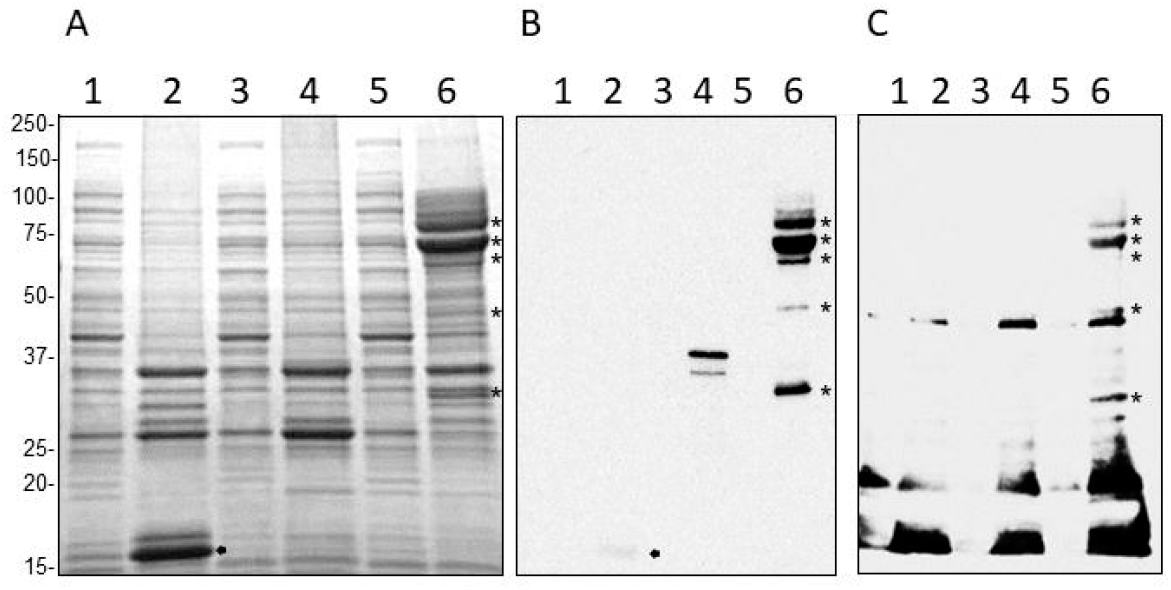
The FhbpA passenger domain binds factor H. **A)** SDS-PAGE gel (10-20% acrylamide gradient) stained with Coomassie brilliant blue R (30 μg protein per lane). **Lanes**: 1 and 2 soluble and insoluble fractions of *E. coli* LDH333 (GroES), respectively; 3 and 4, soluble and insoluble fractions of *E. coli* LDH444 (recombinant autotransporter domain), respectively; 5 and 6, soluble and insoluble fractions of *E. coli* LDH555 (recombinant passenger domain), respectively. **B)** Corresponding far-western (FW) blot probed with a Nickel-HRP probe for the His_6_ tag on recombinant proteins. **C)** Corresponding FW blot probed with human factor H. Passenger domain protein fragments recognized in both FW blots and the stained gel are indicated by stars. Recombinant GroES is arrowed. Molecular weight values from standards are given to the left in kDa.

Identical FW blots were also prepared and probed with human factor H and anti-factor H antibodies. These blots showed that factor H bound to three unrelated *E. coli* proteins of approximately 43.6, 20 and 16 kDa (**Fig. 6C**, lanes 1,2,4 and 6) plus the five FhbpA passenger domain protein bands identified in **Fig. 6B** (lane 6; starred bands). In contrast, a recombinant *B. bacilliformis* GroES control expressed from the same vector (pET-DEST42) or FhbpA’s autotransporter domain (**Fig 6C, lanes 2 and 4**, respectively), did not bind detectable amounts of factor H. These results strongly suggest that the passenger domain of FhbpA is responsible for binding factor H.

### Differential expression of *fhbpA* in response to biologically-relevant environmental conditions and infection of human cells

*B. bacilliformis* must adapt to markedly different environments imposed by the sand fly vector and human host in order to survive. In consideration of the factor H-binding activity of FhbpA, we hypothesized that *fhbpA* gene expression would be greater under “ human-like” versus “ sand fly-like” conditions. To address the hypothesis, we analyzed *fhbpA* expression in response to temperature and pH shifts from the normal cultivation temperature of 30°C and pH of 7.4 to simulate what occurs during transmission between the insect vector and human host. In addition, we examined expression during infection of fresh human blood and low-passage, cultured HUVECs. Results of the RNA-Seq transcriptomic analyses with average *fhbpA* transcripts per million (TPM) are provided in **Table 1**. These results clearly showed enhanced expression of *fhbpA* under conditions that simulated the human host (Pl37, PlBG) or during infection of human blood or vascular endothelial cells (HB37, HBBG, HUVE). The greatest levels of *fhbpA* expression were observed in *B. bacilliformis* RNA samples from infection of HUVEC (HUVE) cells, with an average TPM of 3438 ± 434. In fact, this level of expression was significantly higher relative to all other conditions when analyzed by DESeq2.

**Table 1.** Differential expression of *fhbpA* in response environmental conditions. *B. bacilliformis* cultures (mid-log-phase) were shifted from typical *in vitro* growth conditions to those indicated for 2h, or 24h for HUVEC (HUVE) infections, as described previously [11], then used as source of RNA. Average *fhbpA* transcripts per million (TPM) were determined by RNA-Seq.

| Conditions | Medium | Designation | Simulation | Ave. TPM ± SEM (N)<sup>b</sup> |
| --- | --- | --- | --- | --- |
| pH 7.4,<br>25°C | HIBB plates | PI25 | Sand fly ambient temperature | 106.5 ± 8.5 (2) |
| pH 7.4,<br>30°C | HIBB plates | PI30 | Sand fly ambient temperature | 176.5 ± 32.5 (2) |
| pH 7.4,<br>37°C | HIBB plates | PI37 | Human host | 271.5 ± 17.5 (2) |
| pH 7.4,<br>37°C with<br>blood gas<sup>a</sup> | HIBB plates | PIBG | Human host | 364.7 ± 16.1 (3) |
| pH 6.0,<br>30°C | HIBB liquid | pH06 | Sand fly post-blood meal | 109 ± 2 (2) |
| pH 7.4,<br>30°C | HIBB liquid | pH07 | Human host / sand fly blood<br>meal mid-digestion | 64 ± 5 (2) |
| pH 8.2,<br>30°C | HIBB liquid | pH08 | Sand fly initial blood meal | 69 ± 5 (2) |
| pH 7.4,<br>37°C with<br>blood gas | HUVECs in<br>EGM-Plus<br>medium | HUVE | Human endothelial cell infection | 3,438 ± 434 (2) |
| pH 7.4,<br>37°C | Human<br>blood | HB37 | Human erythrocyte infection | 184.7 ± 14.8 (3) |
| pH 7.4,<br>37°C with<br>blood gas | Human<br>blood | HBBG | Human erythrocyte infection | 223.3 ± 29.5 (3) |

HIBB, Bacto heart infusion blood agar containing 4% defibrinated sheep blood and 2% sheep serum (vol/vol); HUVECs, human umbilical vein endothelial cells; EGM-Plus (Lonza), endothelial cell growth medium containing 2% fetal bovine serum and bovine brain extract.
<sup>a</sup> Blood gas consists of 5% CO<sub>2</sub>, 2.5% O<sub>2</sub>, and 92.5% N<sub>2</sub> at 100% humidity to simulate human blood.
<sup>b</sup> N = number of cDNA libraries constructed and analyzed by RNA-Seq per given condition (biological replicates).

## Discussion

Complement is a cornerstone of humoral innate immunity in vertebrates, as it provides a first line of defense and persistent immune pressure on pathogens. However, despite *Bartonella*’s specialized niche within the mammalian circulatory system, only one study, to date, has examined complement resistance in these bacteria. In that study, Deng et al. correlated resistance of *B. birtlesii* to the presence of the BadA trimeric autotransporter adhesin by demonstrating that sensitivity to complement was significantly increased in a *badA*-knockout strain [24]. In addition, supernatants from wild-type *B. birtlesii* liquid cultures possessed anti-complement activity that could be neutralized by anti-BadA antibodies [24]. In the present study, we describe the complement-resistance phenotype of *B. bacilliformis* with identification and characterization of a serum factor H-binding protein, FhbpA. Interestingly, both this study and the previous report [24] ultimately implicated an autotransporter protein in the complement-resistance phenotype of *Bartonella*. In keeping with these results, previous reports by others have shown that complement resistance can be conferred by surface-exposed, autotransporter proteins of several bacterial pathogens, including the OmpB of *Rickettsia conorii* [5], YadA of *Yersinia enterocolitica* [4, 25], Vag8 and BrkA of *Bordetella pertussis* [7, 26], UspA2 of *Moraxella catarrhalis* [27] and DsrA of *Haemophilus ducreyi* [28, 29].

Autotransporters comprise a large family of outer membrane proteins in Gram-negative bacteria that are involved in virulence. The term “ autotransporter” refers to the ability of these proteins to “ independently” translocate to the outer membrane via type V secretion, as a result of three conserved domains in a single polypeptide chain, including a N-terminal secretory signal peptide for Sec translocon-dependent export across the cytosolic membrane, a C-terminal autotransporter domain, and a passenger domain that is exported to the cell surface through a 12-stranded trans-membrane beta barrel formed by the autotransporter domain (reviewed in [30]; see **Fig. 5**). The domain configuration of FhbpA, together with its Phyre2 structure predictions, suggest that the protein is a type Va or “ classical” autotransporter, as exemplified by the IgA1 protease of *Neisseria gonorrhoeae* [31], the pertactin adhesin of *Bordetella pertussis* [32], and the AIDA-I adhesin of *Escherichia coli* [33]. Our data also suggest that the passenger domain remains associated with the bacterial outer membrane following its secretion (**Fig. 3B**). Moreover, the conserved proteolytic cleavage site between adjacent Arg-Arg residues in the linker regions of secreted serine protease autotransporters [34] is absent in FbhpA’s linker region (see **Fig. 5**). Nevertheless, the possibility exists that some FhbpA passenger domain may be cleaved and released to the medium, as was observed with BadA during growth of *B. birtlesii* in liquid culture [24].

Passenger domains that are not cleaved following export by type Va autotransporters often serve as adhesins for host extracellular matrix proteins and cells [32,33,35]. Thus, our mapping of factor H-binding activity to the passenger domain of FhbpA (**Fig. 6**) was not surprising.

Although FhbpA was not tested for binding to other host proteins, we would not be surprised if the protein was promiscuous regarding host substrate(s), as previously reported for autotransporters from other pathogens that confer serum resistance [4,5,25,27,29].

Molecular Koch’s postulates require that a potential virulence factor be mutagenized in order to gauge the effect of the mutation and to evaluate the determinant’s role in pathogenesis [36]. To this end, we made several attempts to mutagenize *fhbpA* using various suicide vector constructs and our standard protocol for genetic manipulation of *B. bacilliformis* [37]. However, these attempts were unsuccessful, suggesting that the *fhbpA* locus may be essential for viability. In certain respects this result was puzzling, as the complement-resistance phenotype of *B. bacilliformis* in the absence of factor H was considerable (∼49% survival; **Fig. 1**), suggesting that binding of factor H by FhbpA was not essential, and/or the bacterium was protected from complement by mechanisms that didn’t involve factor H. Perhaps additional, undescribed function(s) of FhbpA are essential to viability and precluded mutagenesis. Although not directly demonstrated in this study, we predict that FhbpA provides protection against complement to *B. bacilliformis* by virtue of the ability of bound host factor H to: a) directly bind and neutralize C3b, b) serve as a co-factor for factor I-mediated proteolysis of C3b, and c) accelerate the decay of C3 convertase in the alternative pathway [38].

The widespread occurrence of conserved FhbpA-like passenger domains in the autotransporter proteins of several *Bartonella* species (**Fig. 4**) suggests that binding of serum factor H may be a conserved strategy to enhance complement resistance during infection of various mammalian hosts by members of the genus. *B. schoenbuchensis*, the species with the highest total score in BLASTP searches with the FhbpA passenger domain (**Fig. 4**), can cause bacteremia in ruminants and is possibly transmitted to humans through the bite of infected deer keds [39]. Two other high-scoring hits included the FhbpA-like passenger domains from autotransporters of *B. henselae* and *B. ancashensis*; both recognized human pathogens [1,40].

Transcriptomic analyses of *fhbpA* expression by RNA-Seq suggest that *fhbpA* is an “ infection-specific” gene that is upregulated in response to environmental cues within the human host, including a temperature of 37°C, blood gas (5% CO_2_, 2.5% O_2_ and 92.5% N_2_ at 100% relative humidity) and the appropriate host cells to parasitize (i.e., vascular endothelial cells and erythrocytes) (**Table 1**). Considering the factor H-binding activity of FhbpA, these transcriptomic results are not surprising, since factor H and complement are unique to vertebrates and would only be present in the insect immediately following a blood meal. Nevertheless, it is conceivable that mature FhbpA could play undescribed accessory roles, such as an adhesin, in the context of the sand fly vector [41].

In summary, we have identified four factor H-binding proteins in the membrane fraction of *B. bacilliformis*. One of these was determined to be an autotransporter protein (ABM44634.1) by mass spectrometry, with binding activity conferred by its membrane-associated passenger domain. Widespread occurrence of FhbpA-like passenger domains in other *Bartonella* autotransporters suggests that conserved complement-resistance strategies are employed by the genus. Finally, expression analysis suggests that *fhbpA* expression is upregulated during infection of the human host, especially during the pathogen’s association with vascular endothelial cells.

## Acknowledgments

We are sincerely grateful to Dr. Rich Marconi and Nathaniel O’Bier for technical assistance with FW blots.

## Author Contributions

Conceived and designed the experiments: LH, SW, MM. Performed the experiments: LH, BM, SW, PG, MD, KS. Analyzed the data: LH, BM, SW, MM. Wrote the paper: MM, SW.

