## Supplemental Table 1 for "A human factor H-binding protein of *Bartonella bacilliformis* and potential role in serum resistance"

**S1 Table. Bacterial strains, plasmids and PCR primers used in the study.**

**Strains**

*B. bacilliformis* (*Bb*)

KC583- Neotype strain; ATCC 35685 [1]

*E. coli*

TOP10- Host strain for general cloning (Thermo Fisher)

BL21 Star (DE3)- Host strain for Gateway cloning (Thermo Fisher)

LDH333- *E. coli* BL21 Star (DE3) containing pEXP42-*groES*

LDH444- *E. coli* BL21 Star (DE3) containing pEXP42-AD

LDH555- *E. coli* BL21 Star (DE3) containing pEXP42-PD

**Plasmids**

pCR-XL-TOPO- Cloning vector for long PCR products (Thermo Fisher)

pENTR/D-TOPO- Entry vector for Gateway TOPO cloning (Thermo Fisher)

pET-DEST42- Destination vector for Gateway cloning (Thermo Fisher)

pEXP42-PD- pET-DEST42 with cloned *Bb* *fhbpA* passenger domain

pEXP42-*groES*- pET-DEST42 with cloned *Bb* *groES* gene (BARBAKC583_1173)

pEXP42-AD- pET-DEST42 with cloned *Bb fhbpA* autotransporter domain

**PCR primers**

1. **To clone passenger domain of FhbpA (ABM44634.1; nucleotides 76-2331)**

**Bb1133_PD_CACC_For-** 5’-***CACC*aggaggagcttttgctATG**agtgtccctgtggat-3’

- ***CACC*-** for directional cloning into pENTR/D-Topo
- **aggagg-** RBS
- **agcttttgctATG-** 10 bases upstream of the beginning of the predicted

passenger domain plus a start codon for protein expression.

**Bb1133_PD_Rev-** 5’-CTCACCTTTCTTTTCTACAGGATTTTTCTGC-3’

2. **To clone autotransporter domain of FhbpA (ABM44634.1; nucleotides 2332-3150)**

**Bb1133_TD_CACC_For-** 5’-***CACC*aggagggaaaggtgagATG**caatttgctatctcgc-3’

- ***CACC***- for directional cloning into pENTR/D-Topo
- **aggagg**- RBS
- **gaaaggtgagATG**- 10 bases upstream of start of autotransporter beta

domain plus start codon for protein expression

**Bb1133_TD_Rev-** 5’-AAAAGAAGTTCCTGATGTACCAGATCTTTG-3’

**Citations-**

1. Brenner DJ, O'Connor SP, Hollis DG, Weaver RE, Steigerwalt AG. Molecular characterization and proposal of a neotype strain for *Bartonella bacilliformis*. J Clin Microbiol. 1991 Jul;29(7):1299-302.
